# Expanded Group-Constrained Parcels Enhance Coverage and Reproducibility of the Scene-Selective Cortex

**DOI:** 10.64898/2026.05.16.725633

**Authors:** Yaelan Jung, Hee Kyung Yoon, Rebecca J. Rennert, Daniel D. Dilks

## Abstract

A common approach for investigating high-level visual cortex with functional magnetic resonance imaging (fMRI) is to define regions of interest (ROIs) in individual participants using functional activation clusters and anatomical landmarks. Although highly productive, this approach requires manual decisions about which clusters correspond to specific canonical regions, limiting reproducibility and posing challenges in populations with lower signal-to-noise ratios, such as children. The Group-Constrained Subject-Specific (GSS) approach reduces this subjectivity by using group-level parcels to constrain subject-specific functional ROI definition. However, the original GSS parcel set provides limited coverage of the occipital place area (OPA) and does not include more recently characterized scene-selective regions. Here, we introduce an updated and expanded set of GSS parcels for scene-selective cortex. Using a larger adult sample and dynamic scene stimuli, we generated updated parcels for OPA, parahippocampal place area (PPA), and retrosplenial complex (RSC), and for the first time, delineated a parcel for a newly discovered scene-selective region in the superior parietal lobule (superior place area; SPA). We evaluated these parcels in independent adult and pediatric datasets by testing whether they improve cross-subject coverage while preserving functional selectivity. The updated OPA parcel increased cross-subject coverage relative to the original parcel by Julian and colleagues. Moreover, ROIs defined using the updated parcels showed equal or greater scene selectivity across OPA, PPA, and RSC, indicating improved functional ROI definition without sacrificing specificity. Across scene-selective regions, the updated parcels reliably identified scene-selective cortex and reproduced canonical response profiles and in pediatric data. These parcels provide more complete and reliable coverage of the scene-processing network, supporting objective and reproducible ROI definition across adult and pediatric fMRI datasets.

**Highlights:**

- Expanded group-constrained parcels improve coverage of scene-selective cortex
- Dynamic stimuli yield improved cross-subject overlap for OPA
- New parcel introduced for the scene-selective region in the superior parietal lobule, now called superior place area (SPA)
- Updated parcels reproduce canonical response profiles in adult data
- Parcels reliably identify scene-selective voxels in pediatric datasets

## Introduction

Understanding how the human brain is organized into functionally specialized regions has been a central goal of cognitive neuroscience, with category-selective responses in high-level visual cortex providing a key model system. Functional magnetic resonance imaging (fMRI) studies of category-selective cortex commonly define regions of interest (ROIs) in individual participants by identifying activation clusters that respond more strongly to a category of interest than to a comparison category, such as scenes > objects for scene-selective cortex. These clusters are then assigned to canonical regions based on their anatomical location. This subject-specific functional localization approach has been highly successful for characterizing the functional properties of high-level visual cortex (Dilks & Kanwisher, 2013; Saxe et al., 2006). However, this requires researchers to make manual decisions about which activation clusters correspond to a given ROI. As a result, ROI definition can vary across researchers and datasets, introducing subjectivity and limiting reproducibility. These challenges are especially pronounced in populations with lower signal-to-noise ratios or less reliable individual activation maps, such as children.

Alternative approaches, including atlas-based ROIs or full group-level definitions, avoid manual ROI selection but sacrifice subject-specific functional precision. This can obscure meaningful individual variability in the spatial location and extent of functional regions. To address this tradeoff, Julian et al., (2012) introduced the Group-Constrained Subject-Specific (GSS) approach, building on related approaches for objective functional ROI definition (Fedorenko et al., 2010). In the GSS approach, group-level parcels define constrained search spaces within which subject-specific voxels are selected based on each participant’s own functional activation data. This approach combines the spatial consistency of group-level maps with the functional precision of subject-specific localization, thereby reducing experimenter bias while preserving sensitivity to individual functional organization. As a result, GSS has become a widely used framework for standardizing ROI definition across studies while maintaining subject specificity.

The utility of the GSS approach, however, depends on the quality and completeness of the group-level parcels used to constrain ROI definition. Although the original parcels from Julian et al. (2012) have been widely adopted in studies of category-selective cortex, their performance is not uniform across regions. This issue is particularly relevant for scene-selective cortex. The scene-processing network includes several functionally specialized regions, including the parahippocampal place area (PPA), retrosplenial complex (RSC), and occipital place area (OPA). Among these regions, OPA shows relatively lower cross-subject overlap in the original GSS parcel set, potentially reducing the reliability with which OPA can be identified across individuals. This limitation likely reflects the datasets, stimuli, and known functional regions available at the time of the original release rather than a limitation of the GSS framework itself.

In addition, the known functional architecture of scene-selective cortex has expanded since the original GSS parcels were created. Recent work has identified a scene-selective region in the superior parietal lobule(Yoon et al., 2025, 2026), which we now call the superior place area (SPA), which is functionally linked to visually guided navigation and is often considered alongside OPA as part of a navigation-relevant scene-processing system (Kamps, Lall, et al., 2016; Persichetti & Dilks, 2018; Yoon et al., 2025, 2026). Because SPA was not included in the original Julian 2012 set, the existing GSS framework does not provide complete coverage of currently recognized scene-selective cortex. Together, limited coverage of OPA and the absence of SPA constrain the utility of existing parcels for studies that aim to characterize the full scene-processing network, particularly in datasets where manual ROI definition is difficult or unreliable.

Here, we introduce an updated and expanded set of group-level parcels for scene-selective cortex. To improve OPA coverage, we created a new parcel from a larger adult sample and used dynamic stimuli, which robustly drive responses in OPA and SPA (Kamps, Lall, et al., 2016; Persichetti & Dilks, 2018; Yoon et al., 2025, 2026). We also ensured full parietal coverage during data acquisition, allowing us to generate a new parcel for SPA. In addition, we provide updated parcels for PPA and RSC to create a complete, matched set of scene-selective parcels.

We evaluated these updated parcels in several ways. First, using an independent adult dataset, we compared the updated parcels to the original Julian et al. parcels in terms of parcel size and cross-subject coverage. Second, we tested whether increased coverage came at the cost of functional specificity by comparing scene selectivity in subject-specific ROIs defined using the updated versus original parcels. Third, we assessed whether ROIs defined using the updated parcels exhibit canonical scene-selective response profiles consistent with traditional hand-defined ROIs. Finally, we tested the robustness of the updated parcels in an independent pediatric sample, providing a stringent test of their utility in lower-SNR developmental data. Together, these analyses test whether the updated parcels improve practical ROI definition for scene-selective cortex while preserving the core advantage of the GSS approach: objective, reproducible, and subject-specific functional localization.

## Methods

### Participants

Data were drawn from adult and pediatric participants recruited from the metropolitan Atlanta area across multiple fMRI experiments conducted over several years. The adult sample included 93 participants total. Of these, 72 participants (from three different studies) were used to create the updated group-level parcels, and an independent sample of 21 participants (from another different study) was used to evaluate parcel performance. Across the full adult sample, participants included 60 females, with a mean age of 25.12 years and an age range of 19–41 years. Written informed consent was obtained from all adult participants prior to participation.

The pediatric sample included 97 children: 60 5-year-olds and 37 8-year-olds. The 5-year-old group included 26 females, with a mean age of 5.57 years and an age range of 4.58–6.25 years. The 8-year-old group included 22 females, with a mean age of 8.51 years and an age range of 7.91–9.25 years. Children were recruited through the Emory Child Study Center from the metropolitan Atlanta area across several experiments. Written consent was provided by a parent or legal guardian for all child participants, and verbal assent was additionally obtained from 8-year-old participants. Because the goal of the pediatric analysis was to evaluate parcel performance in lower-SNR developmental data rather than to test age-related differences, data from 5- and 8-year-olds were combined. This approach is also consistent with prior work showing comparable scene selectivity across these age groups in the same regions (Jung et al., 2024; Kamps et al., 2020; Rennert & Dilks, 2025).

All participants had normal or corrected-to-normal vision and no reported history of neurological or psychiatric disorders. All experimental procedures were approved by the Emory University Institutional Review Board.

### Design and Stimuli

Across experiments and age groups, participants completed two functional localizer runs designed to identify scene-selective cortex. The localizer used a blocked design in which participants viewed videos of scenes (Fig. 1A) and objects (Fig. 1B). Each run lasted 180 s and included three blocks per stimulus category. Each stimulus block contained six videos from the same category, with each video presented for 3 s and separated by a 500 ms interstimulus interval, resulting in a total block duration of 21 s. Block order was counterbalanced across runs.

**Figure 1.**
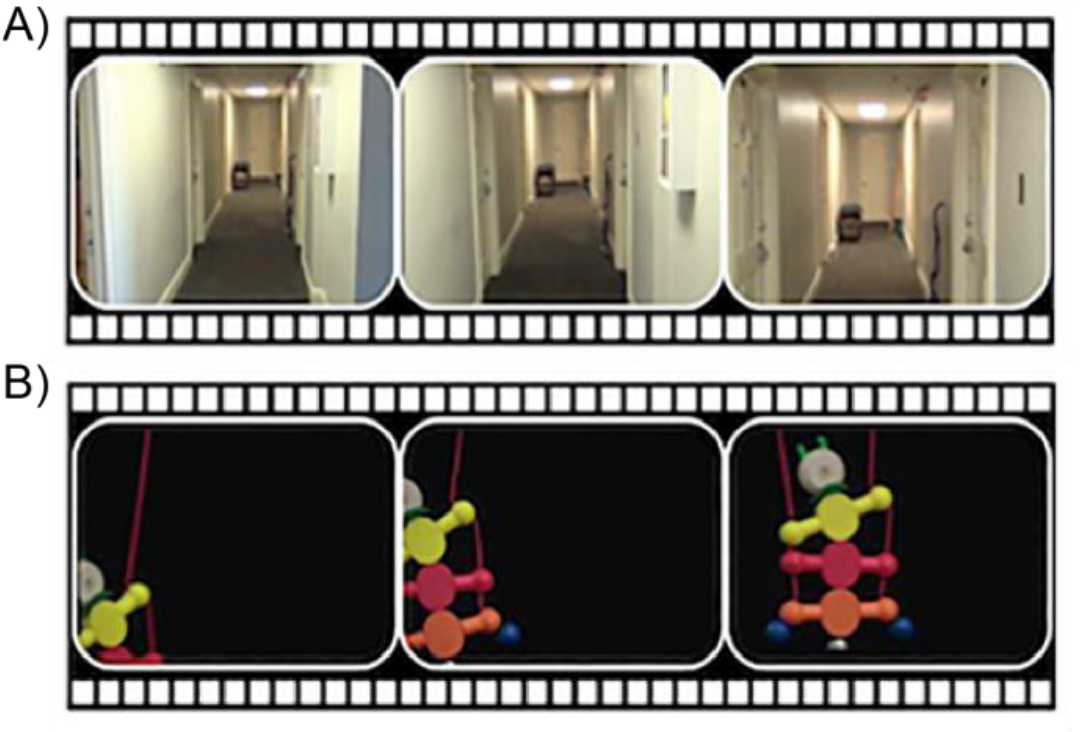
Example dynamic stimuli from Kamps et al. (2020). The conditions included A) Dynamic Scenes, which consisted of 3-s video clips of first-person perspective motion through a scene; B) Dynamic Objects, which consisted of 3-s video clips of moving toys against a black background. Used with permission from Elsevier B.V. Permission granted under a Formal Permission License (FPL).

Each run also included four fixation blocks: one at the beginning of the run, two interleaved between stimulus blocks, and one at the end of the run. Each fixation block lasted 12 s. Participants were asked to watch videos without any task.

### Data acquisition

All scanning was conducted at the Facility for Education and Research in Neuroscience (FERN) at Emory University using either a 3T Siemens Prisma scanner (adult N = 62; children N = 97) or a 3T Siemens Trio scanner (adult N = 31). Functional images were acquired using a 32-channel head matrix coil and a gradient-echo single-shot echo-planar imaging (EPI) sequence. Acquisition parameters were as follows: 28 slices; repetition time (TR) = 2 s; echo time (TE) = 30 ms; voxel size = 1.5 × 1.5 × 2.5 mm; interslice gap = 0.5 mm. Slices were oriented approximately along the anterior commissure–posterior commissure (AC–PC) plane. In addition, high-resolution whole-brain anatomical images were acquired for each participant to support anatomical localization and image registration.

### Initial fMRI data analyses

Preprocessing was performed using AFNI software version 20.3.05. Functional images were registered to each participant’s T1-weighted anatomical image using align_epi_anat.py and corrected for head motion using 3dvolreg. Functional time series were despiked using 3dDespike to reduce transient signal outliers. Spatial smoothing was applied using a Gaussian kernel with a full width at half maximum of 6 mm using 3dmerge. Temporal filtering was applied to remove frequencies above 0.2 Hz. Head-motion parameters relative to the BOLD reference volume were estimated prior to spatial smoothing and temporal filtering.

Following preprocessing, functional data were analyzed using a general linear model implemented in 3dDeconvolve. The model included regressors for the two stimulus conditions, scenes and objects, as well as six motion parameters as nuisance regressors. Stimulus regressors were modeled using BLOCK function. Time points with head motion exceeding 1.5 mm based on framewise displacement were censored from the GLM. Whole-brain contrast maps were computed for scenes > objects and used for parcel creation and ROI evaluation.

### Parcel creation

Updated parcels were created using data from 72 adult participants. These participants were independent from the adult test sample used to evaluate parcel performance. Parcel creation followed the general approach used by Julian et al. (2012), with two key modifications. First, the updated parcels were generated from a larger adult sample (72 subjects versus 30 subjects). Second, the localizer used dynamic scene and object stimuli, which robustly drive responses in OPA and SPA.

For each participant in the parcel-creation sample, a whole-brain scenes > objects contrast map was computed and thresholded at *p* <.001. Individual thresholded contrast maps were transformed into MNI space using 3dQwarp and 3dAllineate in AFNI. Normalized maps were spatially smoothed with a 2 mm Gaussian kernel and averaged across participants to generate a probabilistic overlap map. In this map, each voxel reflected the percentage of participants showing a suprathreshold scenes > objects response at that location.

The probabilistic overlap map was thresholded at 25% to retain voxels showing reliable scene-selective activation across participants. The resulting thresholded map was then segmented using the watershed algorithm implemented in MATLAB version R2021a. This algorithm segments the thresholded overlap map into spatially contiguous parcels centered on local maxima in the group-level activation distribution. Resulting segments were labeled as OPA, PPA, RSC, and SPA based on their anatomical location and correspondence with known scene-selective regions.

The final parcel set included bilateral parcels for OPA, PPA, RSC, and SPA. The SPA parcel was newly introduced because this region was not included in the original Julian 2012 parcel set. Updated PPA and RSC parcels were also included to provide a complete, matched set of scene-selective parcels generated using the same dataset and procedure.

### Evaluation of the parcels

To evaluate whether the updated parcels successfully captured scene-selective voxels corresponding to canonical scene-selective regions, we applied the Group-Constrained Subject-Specific method to independent adult and pediatric datasets. In each participant, the updated parcels were transformed from MNI space into the participant’s native anatomical space and used as constrained search spaces for functional ROI definition.

The GSS procedure was performed using a split-half approach to ensure that voxel selection and response estimation were statistically independent. For each participant and each ROI, voxels within the parcel-defined search space were rank-ordered using data from one localizer run according to the parameter estimates for the scenes > objects contrast. Following Julian et al. (2012), the top 10% of voxels within each parcel were selected to define a subject-specific functional ROI. The response profile of this ROI was then estimated using data from the left-out localizer run. This procedure was repeated with the training and test runs reversed, and response estimates were averaged across the two train-test folds for each participant.

We evaluated parcel performance in three main ways. First, we compared the updated parcels with the original Julian et al. parcels in terms of parcel size and cross-subject coverage. Cross-subject coverage was defined as the percentage of participants with at least one suprathreshold voxel for the scenes > objects contrast within a given parcel. This analysis assessed whether each group-level parcel captured the spatial location of scene-selective activation across individuals.

Second, we compared the functional selectivity of ROIs defined using the updated parcels with ROIs defined using the original Julian et al. parcels. For this analysis, the same split-half GSS procedure was applied separately using each parcel set. For comparisons between updated and Julian 2012 parcels, percent signal change values were analyzed using repeated-measures ANOVAs with parcel version and condition as within-subject factors, conducted separately for each region. The critical test was the parcel version × condition interaction. Significant interactions were followed up with planned paired contrasts comparing scene selectivity, defined as scenes minus objects, between parcel versions.

Third, we assessed whether ROIs defined using the updated parcels reproduced canonical response profiles of scene-selective cortex. Specifically, we tested whether GSS-defined ROIs in OPA, PPA, RSC, and SPA showed greater responses to scenes than to objects. These analyses were conducted first in the independent adult test dataset and then in the pediatric dataset to evaluate parcel robustness in lower-SNR developmental data.

### Traditional hand-defined ROIs

To compare the updated GSS parcels with traditional functional localization procedures, we also identified scene-selective ROIs manually in the independent adult test sample. Hand-defined ROIs were identified in each participant based on the scenes > objects contrast and anatomical location, following standard procedures used in prior studies of scene-selective cortex. ROIs were defined for OPA, PPA, RSC, and SPA where reliable activation was present at *p* < 0.005. Manual ROIs were defined using one run and then tested on the left-out run. Two raters independently created the hand-defined ROIs, and the response from two versions of the manual ROIs were averaged.

Responses in hand-defined ROIs were estimated using an independent run. These response profiles were used to assess whether ROIs defined using the updated parcels produced canonical scene-selective responses consistent with those obtained using traditional manual ROI definition.

### Statistical analyses

Statistical analyses were conducted using R (version 4.5.0). Cross-subject coverage was compared between the updated parcels and the Julian et al. parcels using McNemar’s tests, separately for each region and hemisphere. This test was used because coverage was assessed within the same participants for both parcel versions.

Scene selectivity was analyzed by comparing responses to scenes and objects in independently defined ROIs. For comparisons between updated and Julian et al. parcels, scene selectivity was computed as the response to scenes minus the response to objects for each participant, region, hemisphere, and parcel version. These values were analyzed using repeated-measures ANOVA and paired t-test.

For analyses assessing canonical response profiles, responses to scenes and objects were compared within each GSS-defined ROI using paired-sample t-tests. Effect sizes were reported as Cohen’s d.

### Data and code availability

All parcels are publicly available in both MNI152 nonlinear asymmetric (6th generation) space and MNI152 2009 space to facilitate integration with commonly used neuroimaging pipelines (https://osf.io/zrtxw). We additionally provide example data and a tutorial script demonstrating how to implement the Group-Constrained Subject-Specific (GSS) procedure using these parcels, enabling straightforward adoption of the method across datasets.

## Results

We first describe the creation of the updated parcels for scene-selective cortex. We then evaluate these parcels in an independent adult test dataset by comparing them with the original Julian et al. (2012) parcels in terms of parcel size, cross-subject coverage, and functional selectivity. Finally, we examine the robustness of the updated parcels in a pediatric dataset.

### Creation of the current study parcels

The updated parcels were created using the same general approach as Julian et al. (2012), with two key modifications. First, the present parcels were generated from a larger adult sample than the original parcel set (72 participants vs. 30 participants). Second, we used dynamic stimuli, which robustly drive scene-selective responses in OPA and SPA (Kamps, Lall, et al., 2016; Yoon et al., 2025, 2026). Using individual scenes > objects contrast maps from 72 adults, we first created probabilistic overlap maps in which each voxel reflected the proportion of participants showing suprathreshold scene-selective activation at that location (Fig. 2A). We then thresholded these probabilistic maps and segmented them using a watershed algorithm (Fig. 2B), resulting in an updated set of parcels for scene-selective cortex (see Methods for details). Table 1 summarizes the location, volume, and cross-subject coverage of the updated parcels. These parcels can be transformed into each participant’s native space and used to constrain the search space for identifying subject-specific scene-selective voxels using the GSS approach (Fig. 2C).

**Table 1.** Location, volume, and cross-subject coverage of updated scene-selective parcels.

| Region name | Hemisphere | Coordinate |  |  | Size (mm<sup>3</sup>):<br>Present<br>Study<br>Parcel | Size (mm<sup>3</sup>):<br>Julian 2012 | Percent<br>Subject (%):<br>Present<br>Study Parcel | Percent<br>Subject (%):<br>Julian 2012 |
| --- | --- | --- | --- | --- | --- | --- | --- | --- |
|  |  | x | y | z |  |  |  |  |
| Occipital Place<br>Area (OPA) | LH | 33 | 91 | 13 | 4808 | 2008 | **81.0%** | **66.7%** |
|  | RH | -37 | 85 | 17 | 4840 | 1064 | **85.7%** | **61.9%** |
| Parahippocampal<br>Place area (PPA) | LH | 27 | 41 | -17 | 5448 | 5856 | 95.2% | 90.5% |
|  | RH | -23 | 41 | -17 | 5512 | 4424 | 95.2% | 95.2% |
| Retrosplenial<br>Complex (RSC) | LH | 19 | 57 | 1 | 6032 | 7608 | 90.5% | 90.5% |
|  | RH | -13 | 53 | 1 | 8288 | 8504 | 95.2% | 100% |
| Superior Place<br>Area (SPA) | LH | 17 | 85 | 37 | 4952 | N/A | 90.5% | N/A |
|  | RH | -19 | 77 | 35 | 4064 | N/A | 90.5% | N/A |

**Figure 2.**
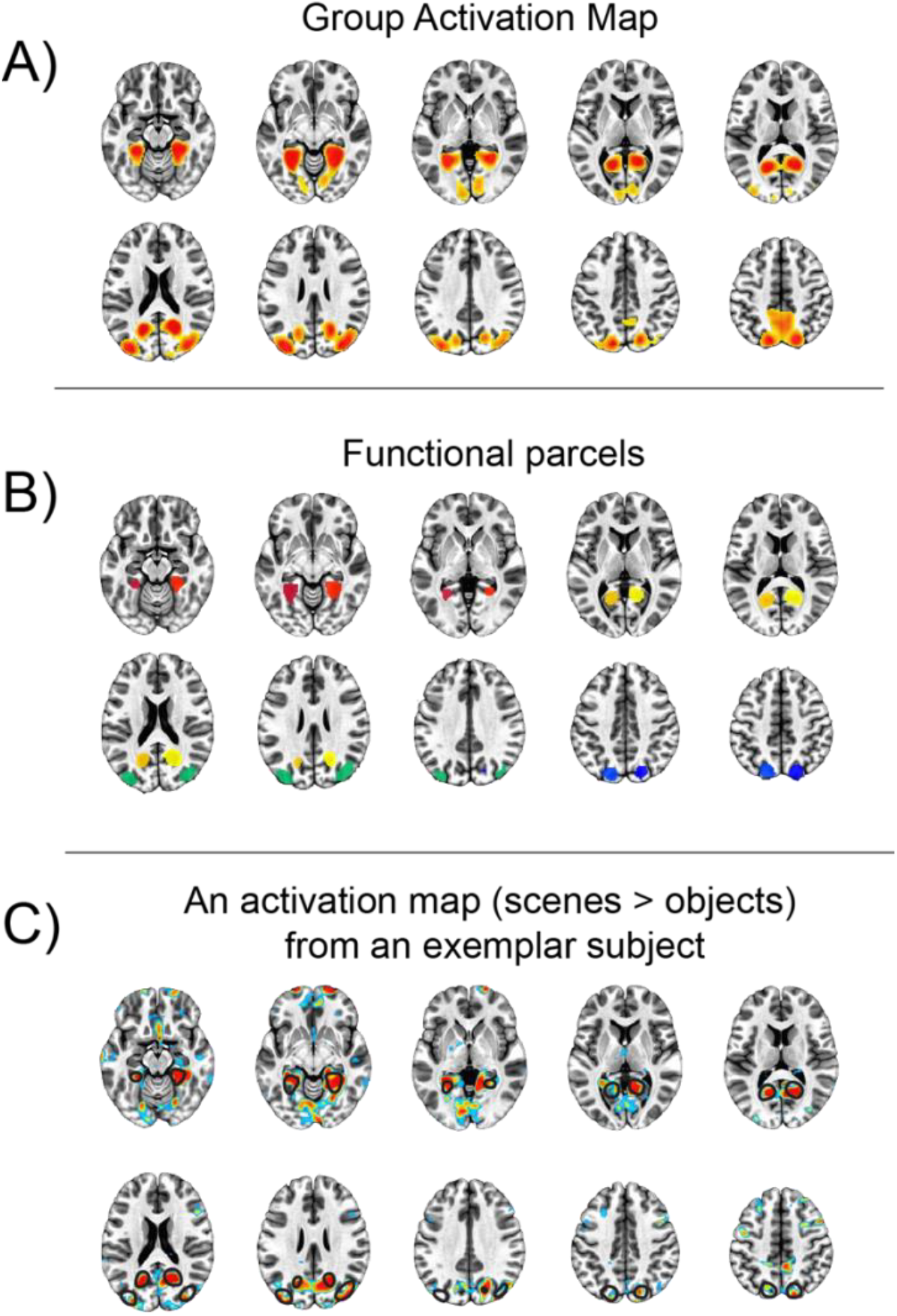
The key steps of creating the group parcel and applying the group-constrained subject-specific method for scene-selective regions. A) Each individual subject scenes > objects activation map is overlaid on top of one another, creating a probabilistic overlap map (n = 72). The color of each voxel corresponds to the percentage of subjects that have activation at that voxel. B) Using a watershed image segmentation algorithm, the overlap map is divided into functional “parcels” following the map’s topography. C) These parcels are then used as spatial constraints to select subject-specific voxels for each region by intersecting each parcel (black outlines) with each individual subjects’ thresholded scenes > objects activation map (*p* < 0.001; following the convention used to define the scene-selective regions).

### Comparison between the two versions of parcels: Coverage

We next quantified cross-subject coverage in an independent adult test dataset (n = 21). Following Julian et al. (2012), coverage was defined as the presence of at least one suprathreshold voxel (*p* < .001) within each parcel for each participant. The updated OPA parcels identified scene-selective voxels in a significantly greater proportion of participants than the Julian et al. parcels in both hemispheres (LH: 81.0% [17/21] vs. 66.7% [14/21], McNemar’s χ^2^(1) = 5.556, *p* = .018; RH: 85.7% [18/21] vs. 61.9% [13/21], χ^2^(1) = 6.250, *p* = .012), indicating improved cross-subject coverage (also see Fig. 3).

**Figure 3.**
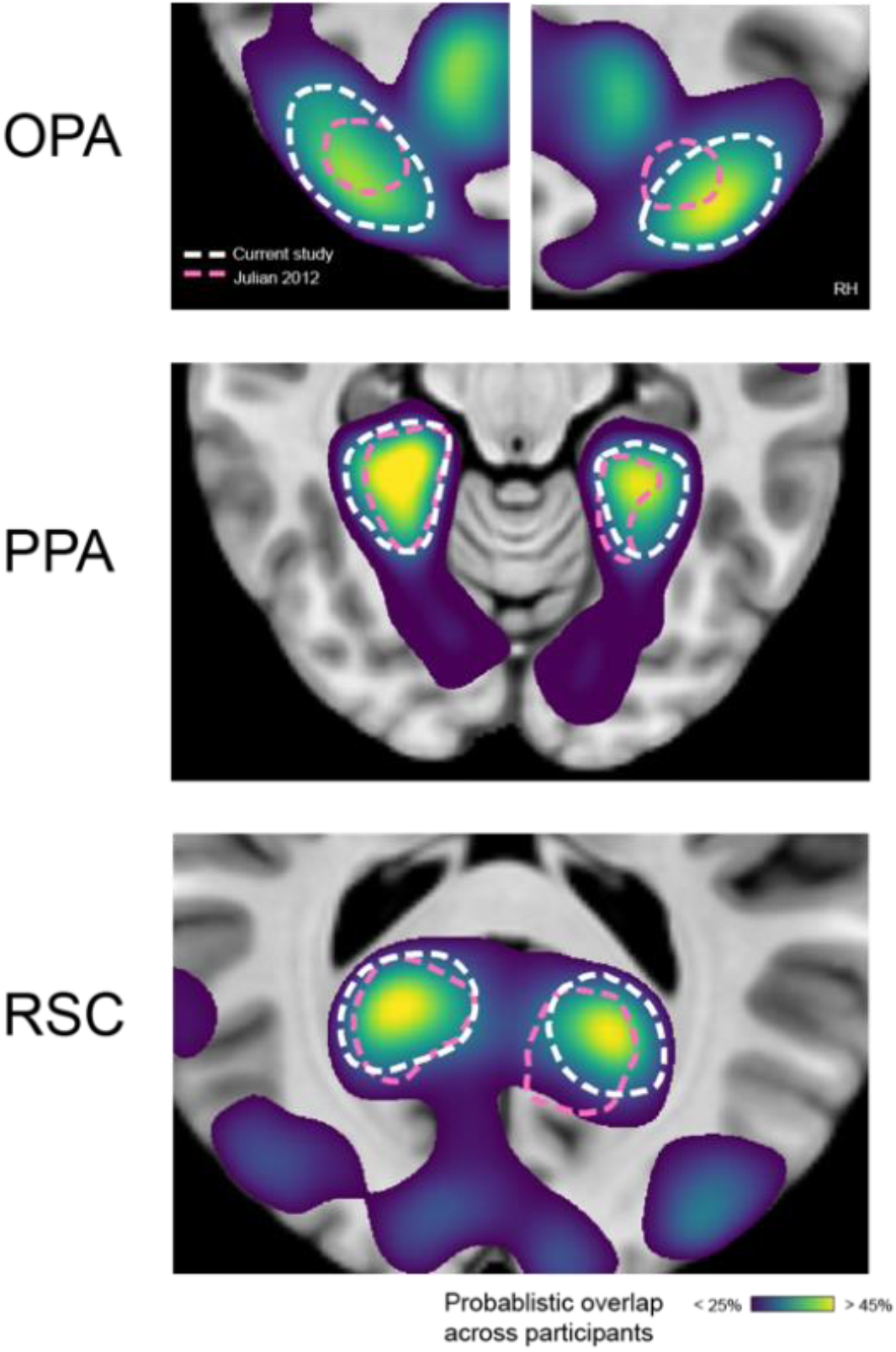
Probabilistic overlap maps for scene-selective regions with parcel outlines from the current study and Julian et al. (2012). The color maps indicate the proportion of participants in the parcel-creation sample showing suprathreshold activation for the contrast *scenes > objects* at each voxel. White dashed outlines indicate parcels from the current study, and pink dashed outlines indicate the original Julian et al. (2012) parcels. Shown are bilateral parcels for OPA, PPA, and RSC.

Coverage rates were similar across parcel versions for PPA and RSC. For PPA, coverage was 95.2% [20/21] for the updated parcel versus 90.5% [19/21] for the Julian et al. parcel in the LH, χ^2^(1) = 0.333, *p* = .564, and 95.2% [20/21] for both parcel versions in the RH, χ^2^(1) = 0, *p* = 1. For RSC, coverage was 90.5% [19/21] for both parcel versions in the LH, χ^2^(1) = 0, *p* = 1, and 95.2% [20/21] for the updated parcel versus 100% [21/21] for the Julian et al. parcel in the RH, χ^2^(1) = 0.333, *p* = .564. Thus, the updated parcels selectively improved cross-subject coverage for OPA, while maintaining high coverage for PPA and RSC.

### Comparison between the two versions of parcels: Scene Selectivity

Greater cross-subject coverage does not necessarily imply improved functional ROI definition. A larger parcel could identify scene-selective activation in more participants while also including additional noise, thereby reducing scene selectivity. To test whether the updated parcels indeed captured additional meaningful scene-selective signal rather than noise, we compared responses in subject-specific ROIs defined using the updated parcels and the Julian 2012 parcels, using independent data from the left-out localizer run (n = 21).

For each region, we conducted a repeated-measures ANOVA with parcel version (updated parcels, Julian 2012 parcels) and condition (scene, object) as within-subject factors. The critical test was the parcel version × condition interaction, which asked whether the magnitude of scene selectivity (scene – object) differed between ROIs defined using the updated versus Julian 2012 parcels. First, in OPA, this parcel version by condition interaction was significant, *F*(1,20) = 9.321, *p* = 0.003, ηp^2^ = 0.318. Planned contrast tests revealed that the updated OPA parcel yielded stronger scene selectivity (scene – object) than the original Julian 2012 parcel (updated: 0.67 ± 0.37; Julian et al.: 0.43 ± 0.29; *p* = 0.018), while both versions of parcels detected significantly greater response to scenes over objects (updated OPA parcel: *p* < 0.001; Julian 2012 parcel: *p* < 0.001; planned simple effect tests) (Fig. 4A). Thus, the expanded OPA parcel yielded a greater magnitude of scene-selective responses in OPA, likely due to its greater cross-subject coverage (Table 1; Fig. 3).

**Figure 4.**
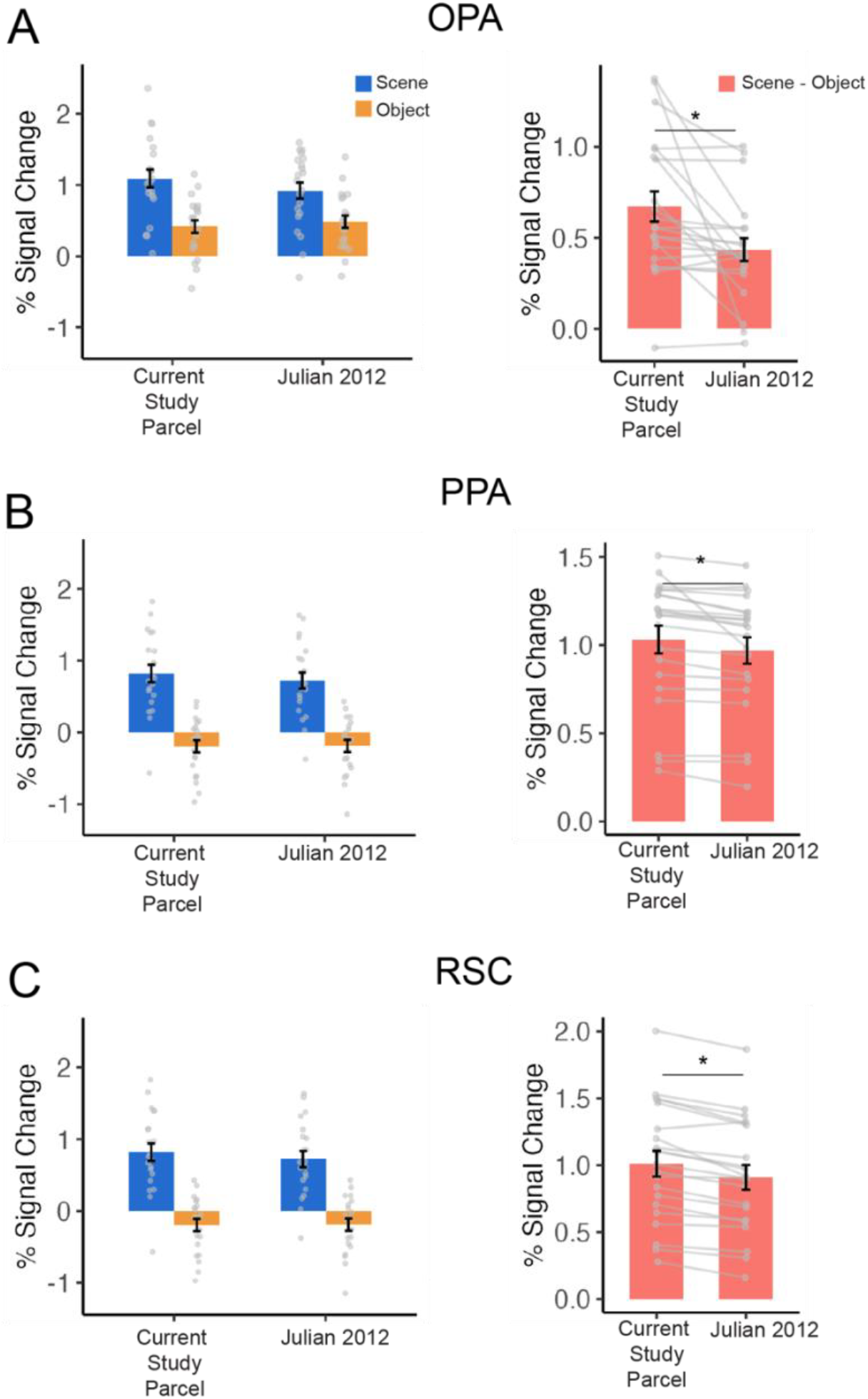
The mean percent signal change for scene (blue) and object (orange) conditions (left panel), averaged across functionally identified voxels for OPA (A), PPA (B), RSC (C), and the difference between scene and object (i.e., scene selectivity) across three regions (right panel). Each bar presents the average across group, error bars represent standard error, and dots represent individual data point. * *p* < 0.05.

Unexpectedly, the updated parcels also yielded stronger scene selectivity in PPA and RSC. In PPA, the parcel version × condition interaction was significant (Fig. 4B), *F*(1,20) = 8.206, *p* = 0.01, ηp^2^ = 0.291. Again, planned contrast tests revealed greater scene selectivity for PPA defined using the updated parcel than the Julian 2012 parcel (updated: 1.02 ± 0.37; Julian 2012: 0.96 ± 0.34; *p* = 0.009), while planned simple-effect tests reveal that both parcels successfully detected scene selectivity (the updated parcel: *p* < 0.001; the Julian 2012 parcel, *p* < 0.001).

Similarly, in RSC, the parcel version × condition interaction was significant (Fig. 4C), *F*(1,20) = 47.322, *p* < 0.001, ηp^2^ = 0.703, with planned contrast tests revealing stronger scene selectivity for ROIs defined using the updated parcel than the Julian 2012 parcel (updated: 1.01 ± 0.44; Julian 2012: 0.91 ± 0.42; *p* < 0.001). Again, planned simple-effects tests confirmed significant scene selectivity for both the updated parcel (*p* < 0.001) and the Julian 2012 parcel (*p* < 0.001). Together, these results indicate that ROIs defined using the updated parcels show stronger scene-selective response profiles than ROIs defined using the Julian 2012 parcels.

### Updated parcels reproduce canonical response profiles observed with manual ROI definition

Having shown that the updated parcels improve GSS-based ROI definition relative to the original Julian 2012 parcels, we next confirm that ROIs defined using the updated parcels along with the GSS approach exhibited response profiles consistent with those obtained using traditional manual ROI definition. This analysis was intended as a qualitative validation of the updated parcels to ensure it can replace the traditional manual approach when needed.

Across OPA, PPA, RSC, GSS-defined ROIs showed the expected response profile (Fig. 5A-5C), with stronger responses to scenes than objects. These response profiles closely resembled those observed in manually defined ROIs. For completeness, GSS-defined ROIs showed significant scene selectivity in all four regions, all *p*s < .001, just as manually defined ROIs, all *p*s < 0.001.

**Figure 5.**
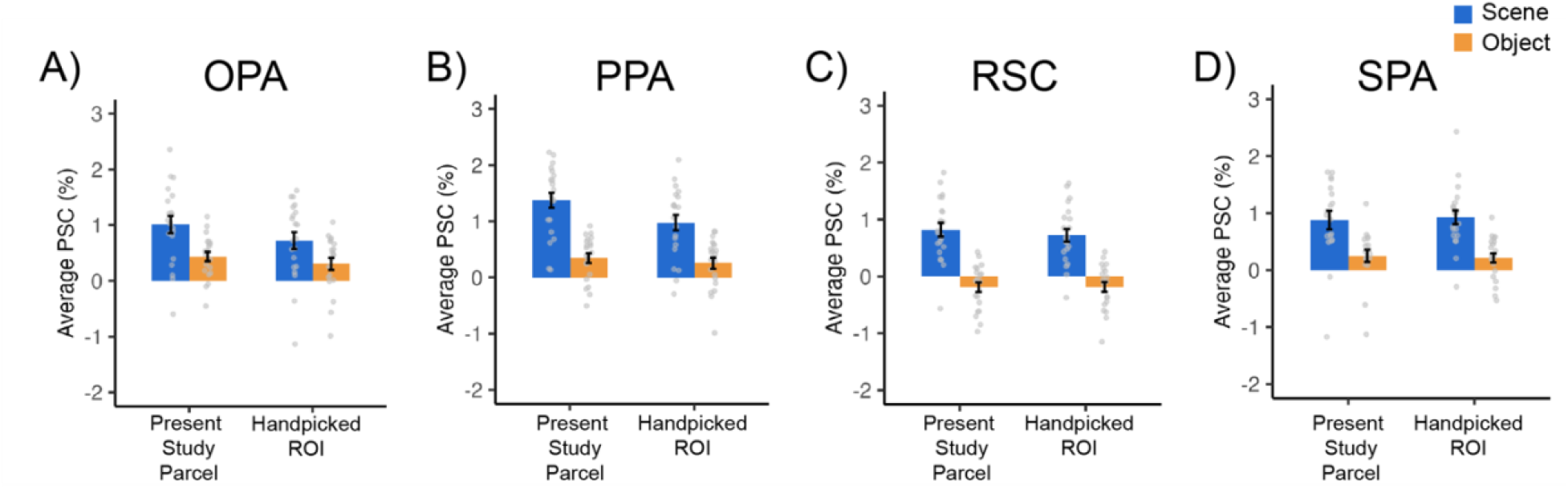
Response profile of the four scene-selective regions when functional ROIs were identified with GSS method and the present study parcel versus the traditional handpicking approach. Across all four scene-selective regions, greater response to scenes over object (scene selectivity) is captured consistently across two methods.

Importantly, we found the same pattern in the newly discovered scene-selective region, SPA (Fig. 5D). The scene selectivity detected using the SPA parcel resembled the pattern of responses observed in manually defined SPA, and both approaches successfully detected greater responses to scene over objects (all *p*s < 0.001).

Taken together, the updated parcels provide valid search spaces for identifying subject-specific scene-selective ROIs using the GSS approach, including both canonical scene-selective regions and the more recently characterized SPA.

### Evaluation of the current parcels in the pediatric sample

We next evaluated whether ROIs defined using the updated parcels exhibit scene-selective responses in a pediatric dataset (N = 97; Fig. 6). The updated parcels were transformed to each participant’s native space and used to define functional ROIs using the GSS method. Across all three scene-selective regions (OPA, PPA, and RSC), we conducted separate repeated-measures ANOVAs for each region with parcel version (the current parcel, Julian 2012 parcel) and condition (scene, object) as within-subject factors to test if the updated parcel better detects scene selectivity than the Julian 2012 parcels.

**Figure 6.**
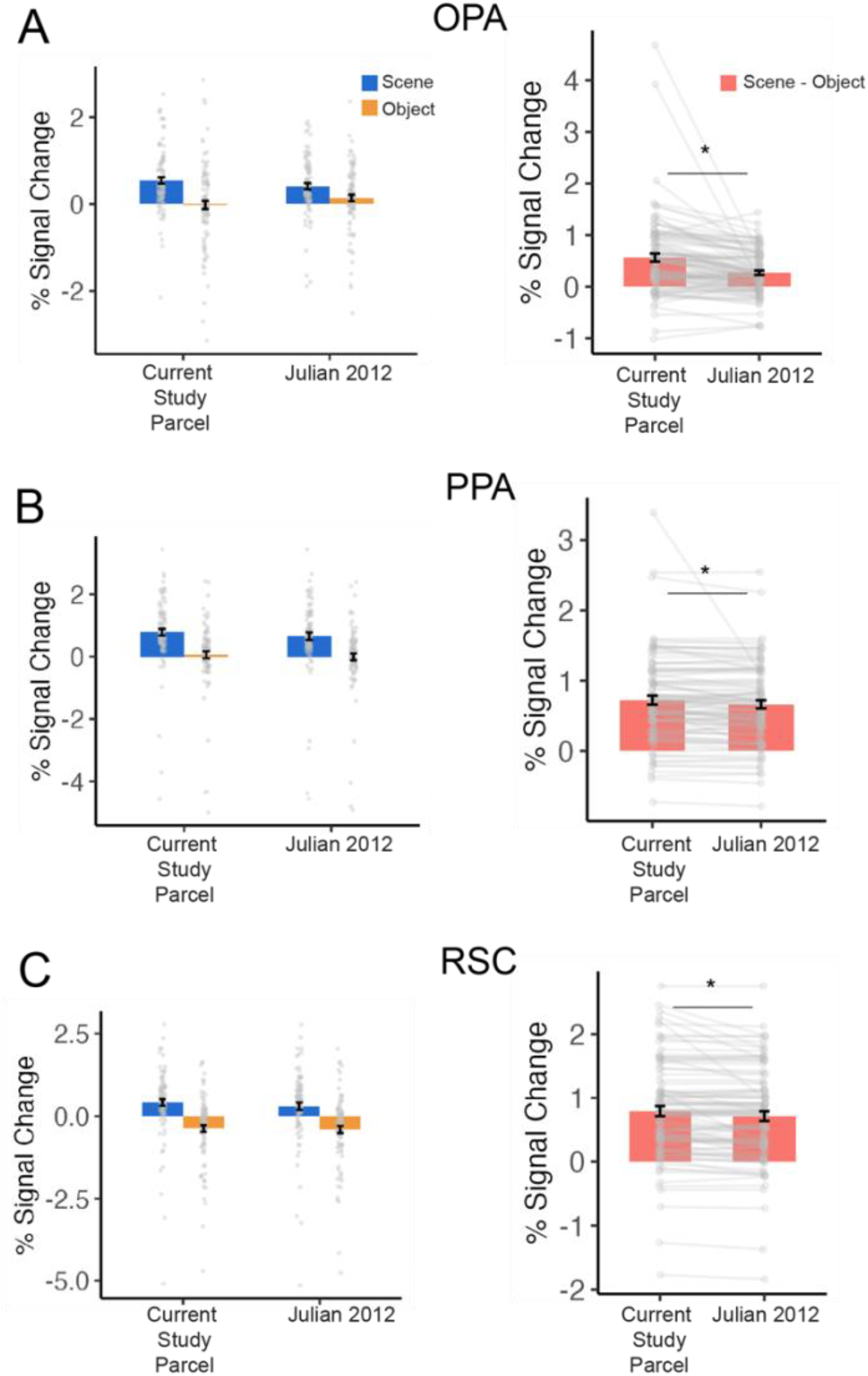
The mean percent signal change for scene (blue) and object (orange) conditions (left panel) in children, averaged across functionally identified voxels for OPA (A), PPA (B), RSC (C), and the difference between scene and object (i.e., scene selectivity) across three regions (right panel). Each bar presents the average across group, error bars represent standard error, and dots represent individual data point. *p* < 0.05.

In OPA, as predicted, we found significant interaction between parcel version and condition, *F*(1,96) = 22.823, *p* < 0.001, η_p_^2^ = 0.192, with greater scene selectivity observed in the updated parcel than the Julian 2012 parcel (*p* < 0.001; planned contrast tests). Like in adults, while unexpected, we found the same pattern in both PPA and RSC (PPA: *F*(1,96) = 5.393, *p* = 0.02, η_p_^2^ = 0.053; RSC: *F*(1,96) = 15.436, *p* < 0.001, η_p_^2^ = 0.139), with greater scene selectivity detected with the updated parcel compared to the Julian 2012 parcel (PPA: *p* < 0.005; RSC: *p* < 0.001; planned contrast tests).

Next, to further examine whether the updated parcels improve ROI definition in pediatric data, we compared the number of children in whom a scene-selective ROI could be successfully identified with each parcel version. Successful ROI identification was defined as greater responses to scenes than objects within the GSS-defined ROI for a given participant and region. For instance, if the average response of a GSS-defined ROI is not greater for scene compared to object, we consider that a given scene-selective ROI is not identified for a given participant. As predicted, this advantage was most evident in OPA: scene-selective OPA ROIs were identified in [80/97; 82.5%] children using the updated parcel, compared with [68/97; 70.1%] children using the Julian 2012 parcel, a significant difference by McNemar’s test, χ^2^(1) = 6.7222, *p* = 0.009. By contrast, although the updated parcels yielded stronger scene-selective responses in PPA and RSC, they did not significantly increase the number of children in whom scene-selective ROIs were detected (PPA: updated parcel = [86/97; 88.7%], Julian 2012 parcel = [86/97; 88.7%], χ^2^(1) = 0, *p* = 1; RSC: updated parcel = [87/97; 89.7%], Julian 2012 parcel = [85/97; 87.6%], χ^2^(1) = 0.5, *p* = 0.4795). Thus, the updated parcels improved both the magnitude and detectability of scene-selective responses in OPA, while producing stronger response profiles in PPA and RSC without significantly changing ROI detection rates.

Finally, we also tested if scene-selective responses can be detected in SPA by transforming the SPA parcel to each participant’s native space and using the GSS approach. A one-way ANOVA with condition (scene, object) as a within-subject factor revealed significant effect of condition, *F*(1,96) = 49.828, *p* < 0.001, η_p_^2^ = 0.342, suggesting that like in adults, scene selectivity in SPA can be defined in pediatric data using our parcel (Fig. 7).

**Figure 7.**
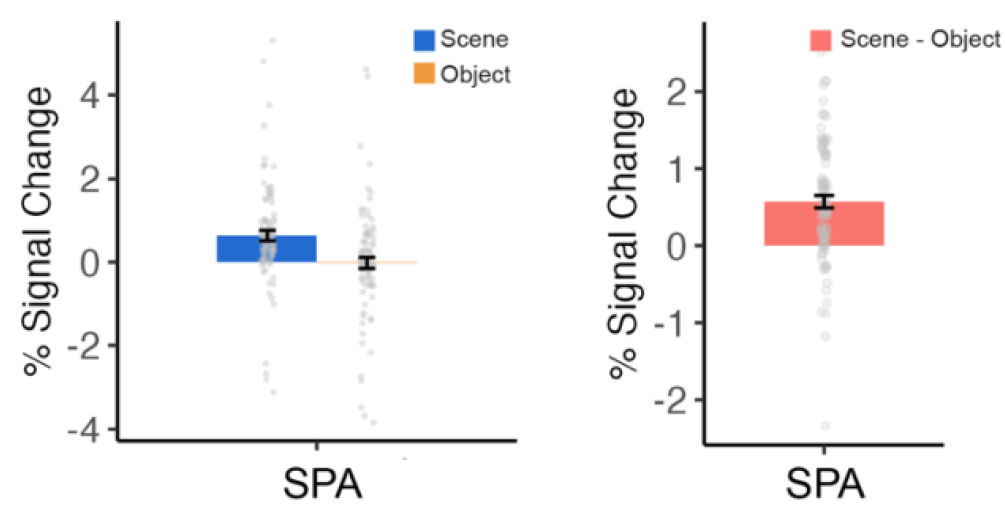
The mean percent signal change for scene (blue) and object (orange) conditions (left panel) in SPL in children, and the difference between scene and object (i.e., scene selectivity) in SPA (right panel). Each bar presents the average across group, error bars represent standard error, and dots represent individual data point.

Taken together, these findings demonstrate that the updated parcels can be applied to datasets with greater anatomical variability, supporting their robustness across developmental populations.

## Discussion

The present study introduces an updated set of group-level parcels for the scene-selective regions, including an improved parcel for OPA and a new parcel for the recently characterized scene-selective region, SPA (Yoon et al., 2025, 2026). The updated OPA parcel was derived using dynamic (movie) scene stimuli, which elicit stronger responses in OPA than static images (Kamps, Lall, et al., 2016; Yoon et al., 2025), and from a larger sample than that used in Julian et al. (2012), with the goal of increasing cross-subject coverage and reproducibility. Using an independent adult dataset, we then evaluated how the updated parcels compare to the original Julian 2012 parcels and how effectively they identify scene-selective voxels when used within the Group-Constrained Subject-Specific (GSS) framework. We found that the new OPA parcel is larger and shows greater cross-subject overlap than the original parcel, while parcels for PPA and RSC remain largely comparable to those from the original release. Importantly, the scene-selective regions defined using the updated parcels exhibited response profiles similar to those obtained with manually defined ROIs in adult data and reliably captured scene selectivity when applied to pediatric datasets. Together, these findings indicate that the updated parcels provide a robust and generalizable tool for defining scene-selective regions.

The updated OPA parcel is particularly relevant for ongoing research investigating how the human brain supports visually guided navigation. A growing body of research has examined the types of information represented in OPA (Bonner & Epstein, 2017; Jones et al., 2023; Kamps et al., 2025; Kamps, Julian, et al., 2016; Kamps, Lall, et al., 2016; Park & Park, 2020; Persichetti & Dilks, 2016) and how this region contributes to navigation-relevant processing (Julian et al., 2016; Persichetti & Dilks, 2018), as well as how this system develops across childhood (Dilks et al., 2023; Jung et al., 2024; Kamps et al., 2020; Rennert & Dilks, 2025). Compared to other scene-selective regions such as PPA and RSC, OPA often exhibits greater anatomical variability across individuals, which can make it challenging to define consistently—especially when activation clusters are weak or ambiguous. By increasing cross-subject coverage, the updated parcel may facilitate more reliable identification of OPA across studies and populations, thereby improving measurement precision and comparability across datasets.

The present release also provides a group-level parcel for SPA, enabling more consistent investigation of this recently identified scene-selective region (Yoon et al., 2025, 2026). This parcel can serve as an anatomical and functional reference for studies seeking to localize SPA and can be used directly to define subject-specific SPA ROIs with the GSS method. A dedicated SPA parcel is useful because SPA may be anatomically close to, but distinct from, other recently reported scene-responsive regions. For example, PIGS (Kennedy et al., 2024) was defined using a related but distinct contrasts (scenes > faces or scenes > non-scenes), and appears to be anatomically distinct from SPA (Yoon et al., 2025), and lacks true scene selectivity by responding equally to static objects and static scenes (Yoon et al., 2026). Thus, the SPA parcel provides a more precise reference for studies specifically targeting SPA and helps distinguish it from nearby regions identified using different functional criteria. As with the updated parcels for canonical scene-selective regions, this may be especially useful in populations with noisier data, such as pediatric samples, where objective constraints can improve the reliability of functional ROI definition.

Finally, although the largest expected improvement was in OPA, the updated parcels also yielded modestly stronger scene selectivity in PPA and RSC. This suggests that the updated parcels may capture somewhat more selective voxels even in regions that were already well characterized by the Julian et al. (2012) parcels. However, this improvement should be interpreted cautiously: unlike in OPA, the updated PPA and RSC parcels did not significantly increase the number of children in whom scene-selective ROIs could be identified. Thus, for PPA and RSC, the advantage of the updated parcels appears to reflect a modest strengthening of response profiles rather than a major improvement in ROI detectability. This effect may reflect the use of a larger parcel-creation sample, dynamic stimuli, or the creation of a matched parcel set using a consistent procedure. However, because the current study was not designed to isolate which feature of the updated procedure drove these improvements, this interpretation remains tentative.

In summary, the updated parcel set extends the GSS framework by improving coverage of OPA and incorporating SPA, while maintaining response profiles comparable to traditional hand-defined functional ROIs. By providing validated, publicly available parcels, this resource supports more objective, reproducible, and consistent definition of scene-selective cortex across studies and populations.

## Acknowledgments

This work was supported by a grant from the National Eye Institute (R01 EY29724 to D.D.D).

